# Determining the link between alpha-gal-containing antigens in North American ticks and red meat allergy

**DOI:** 10.1101/505776

**Authors:** Gary Crispell, Scott P. Commins, Stephanie A. Archer-Hartman, Shailesh Choudhary, Guha Dharmarajan, Parastoo Azadi, Shahid Karim

**Affiliations:** Department of Cell and Molecular Biology, School of Biological, Environment and Earth Sciences, The University of Southern Mississippi, Hattiesburg, MS 39406, USA; Department of Medicine & Pediatrics, University of North Carolina, Chapel Hill, NC 27599-7280, USA; Complex Carbohydrate Research Center, University of Georgia, Athens, Georgia 30602-4712, USA; Savannah River Ecology Laboratory, University of Georgia, Aiken, South Carolina 29808, USA

**Keywords:** Alpha-gal, red meat allergy, ticks, saliva, salivary glands, glycans

## Abstract

Development of specific IgE antibodies to the oligosaccharide galactose-α-1, 3-galactose (α-gal) following tick bites has been shown to be the source of red meat allergy. In this study, we investigated the presence of α-gal in four tick species: the lone-star tick (*Amblyomma americanum*), the Gulf-Coast tick (*Amblyomma maculatum*), the American dog tick (*Dermacentor variabilis*), and the black-legged tick (*Ixodes scapularis*) by using a combination of immunoproteome, carbohydrate analysis, and basophil activation approaches. Using anti-α-gal antibodies, α-gal was identified in the salivary glands of both *Am. americanum* and *Ix. scapularis*, while *Am. maculatum* and *De. variabilis* appeared to lack the sugar. PNGase F treatment confirmed the deglycosylation of N-linked α-gal-containing proteins in tick salivary glands. Immunolocalization of α-gal moieties to the salivary secretory vesicles of the salivary acini also confirmed the secretory nature of α-gal-containing antigens in ticks. *Am. americanum* ticks were fed human blood (lacks α-gal) using an artificial membrane feeding system to determine the source of α-gal. N-linked glycan analysis revealed that *Am. americanum* and *Ix. scapularis* have α-gal in their saliva and salivary glands, but *Am. maculatum* contains no detectable quantity. Salivary samples from *Am. americanum* and *Ix. scapularis* stimulated activation of basophils primed with plasma from α-gal allergic subjects. Together, our data support the idea that bites from certain tick species may specifically create a risk for the development of α-gal-specific IgE and hypersensitivity reactions in humans. Alpha-Gal syndrome challenges the current food allergy paradigm and broadens opportunities for future research.

## Introduction

Food allergies are a growing food safety and public health concern, and according to the Centers for Disease Control and Prevention, food allergies are estimated to affect 4%–6% of children, and 4% of adults in the United States^1^. While food allergy symptoms are most common in infants, children, and adolescents, they can appear at any age. Additionally, the symptoms and severity of allergic reactions to specific foods can vary between individuals and can change within a person over time. Most food-related symptoms occur between a few minutes to 2 h after ingestion. Acute, potentially fatal, anaphylactic reactions generally are manifested within minutes of exposure to the trigger food^2^. However, delayed hypersensitivity reactions to foods (e.g., eczema) are not uncommon, and generally develop several hours after allergen exposure^3^. While food allergies generally are associated with immune responses to specific proteins, a novel IgE antibody response to the oligosaccharide epitope galactose-alpha-1,3-galactose (alpha-gal or α-gal) found in mammalian food products (e.g., beef and pork) has been reported^4^. This delayed hypersensitivity, termed red meat allergy or alpha-gal syndrome (AGS), appears to develop at any age, often with several decades of clear immunologic tolerance to mammalian meat.

Although AGS has been identified worldwide, in the United States a growing body of research suggests that bites from the lone-star tick (*Amblyomma americanum*) give rise to α-gal-specific IgE (sIgE)^5^, and this is an unusual allergic reaction to mammalian meat products^6^. In some instances, tick bites have been specifically indicated as the likely mechanism of red meat allergy^7,8^. However, many physicians remain unaware of this growing problem, and patients can often be misdiagnosed with an idiopathic hypersensitivity reaction^9^. Outside of the US, other tick species have been identified that may also be involved in the development of AGS including *Ixodes holocyclus* in Australia^10^, *Ixodes ricinus* in Europe^11^, *Haemaphysalis longicornis* in Japan^12^, and *Amblyomma sculptum* in Brazil^13^. As lone-star ticks have spread from the Southwest to the East Coast of the US, the reported number of individuals suffering allergic reactions after eating red meat has been increasing. The carbohydrate, galactose-α-1,3-galactose (α-gal), can be found in beef, lamb, pork, and food products derived from all mammals other than catarrhine primates (apes and humans), and digestion releases the antigenic glycans resulting in a delayed-type allergic response^14^. Normally, α-gal found in red meat poses no risk to humans, but after attachment of the lone-star tick to the host, it is possible that α-gal-containing antigens from the tick delivered into the host’s skin trigger an α-gal-directed IgE response. Because all immunocompetent humans develop IgM, IgG, IgA, and IgD responses to α-gal^15^, an alternative explanation is that the bite of *Am. americanum* induces a Th2 response in the host, which skews the human immune system to begin producing an IgE class antibody response to α-gal.

Surprisingly, continued exposure to tick bites seems to augment the already existing sIgE antibody response. The important questions we asked in this study were how do ticks induce IgE responses, and why is the response it directed specifically to α-gal? Our findings revealed that α-gal was present in both *Am. americanum* and *Ix. scapularis* and identified the tick antigens that could be potentially associated with the development of the α-gal-directed IgE immune response in humans.

## Materials and Methods

### Ethics statement

All animal experiments were conducted in strict accordance with the recommendations in the Guide for the Care and Use of Laboratory Animals of the National Institutes of Health, USA. The protocol for tick blood feeding on sheep was approved by the Institutional Animal Care and Use Committee of the University of Southern Mississippi (protocol # 15101501). All efforts were made to minimize animal suffering.

### Materials

All common laboratory supplies and chemicals were purchased from Sigma-Aldrich (St. Louis, MO, USA), Fisher Scientific (Grand Island, NY, USA), or Bio-Rad (Hercules, CA, USA) unless otherwise specified.

### Ticks and other animals

The lone-star tick (*Amblyomma americanum*), Gulf-Coast tick (*Amblyomma maculatum*), American dog tick (*Dermacentor variabili*s), and the black-legged tick (*Ixodes scapularis*) were maintained at the University of Southern Mississippi according to established methods^16^. Unfed adult ticks were purchased from Oklahoma State University’s tick rearing facility (Stillwater, OK, USA). Adult ticks were kept at room temperature with approximately 90% relative humidity under a photoperiod of 14 h of light and 10 h of darkness before infestation on sheep. Adult ticks were blood-fed on sheep and removed at intervals between 1 and 11 days, depending upon the experimental protocol.

### Artificial membrane feeding

A silicone membrane-based artificial feeding system was used to feed ticks on human blood as previously described^17^ with minor modifications. Membranes were constructed by using a silicone oil solution to coat lens paper that was allowed to cure for at least 48 h. After curing, a fiberglass mesh was attached between the surfaces of the membrane and an acrylic chamber using silicone adhesive. After curing for 24 h, the seal was verified by immersing the chamber in 70% ethanol for 20 min^18^. Ten virgin females and four male *Am. americanum* ticks were placed into each feeding chamber with sheep hair to compensate for the host odor. Defibrinated whole human blood (Bioreclamation IVT, Westbury, NY, USA) was used for the artificial membrane feeding of ticks in this study. Human blood was stored at 4°C, and 3–4 mL aliquots were warmed to 37°C and added to a single well of a 6-well plate. The feeding chamber was placed into the well so that the membrane came into direct contact with the human blood. Each chamber was blocked with a cotton stopper to isolate the ticks to that area. To maintain an optimal feeding temperature, the system was placed in a 37°C incubator. The blood was replaced at 12-h intervals, and the membranes and six-well plates were rinsed with a solution of 1× PBS containing 5% penicillin/streptomycin. The chambers were monitored daily for changes in attachment rate, size, feeding success, and mortality.

### Tick tissue dissections and saliva collection

The adult female ticks that were blood-fed were dissected within hours of removal and collection from the sheep as described previously^19^. Tick tissues were dissected and washed in M-199 buffer^20,21^. Tissues were stored at −80°C in 0.15 M Tris-HCl, pH 8.0, containing 0.3 M NaCl, 10% glycerol, and 1% protease inhibitor cocktail (Amresco, Solon, OH, USA). Tick saliva was collected by inducing partially-blood-fed female *Am. americanum* to salivate into capillary tubes using the modified pilocarpine induction method as described previously^22,23^. The saliva was stored immediately at −80°C until subsequent western blot analysis.

### Protein extraction

Proteins were solubilized from dissected salivary glands and midgut tissues in a protein extraction buffer consisting of 0.5 M Tris-HCl, pH 8.0, 0.3 M NaCl, and 10% glycerol, and were then treated with 1% HALT protease inhibitor cocktail. The tissues were crushed using pestles and sonicated using a Bioruptor Pico (Diagenode, Denville, NJ, USA) sonication device for 10 full cycles of 30 s pulse/30 s rest at 4°C. Homogenates were centrifuged at 5000 x g for 10 min at 4°C and the supernatants were collected. Protein concentrations were estimated using the Bradford method^24^, and protein was stored at −80°C.

### SDS-PAGE and western blotting

Extracted proteins from the midguts (15 µg), salivary glands (15 µg), and saliva (10 µg) were fractionated on a Mini-PROTEAN TGX Any kD, 7.5%, or 4%–20% gels (Bio-Rad) using SDS-PAGE and were then transferred onto nitrocellulose membrane in a Transblot cell (Bio-Rad). The transfer buffer comprised 25 mM Tris-HCl and 192 mM glycine in 20% methanol. Nonspecific protein binding sites were blocked with 5% BSA in a TBS and Tween-20 solution, and the membranes were incubated with α-galactose (M86) monoclonal IgM antibodies (Enzo Life Sciences, Farmingdale, NY, USA) using an iBind western device (Life Technologies, Camarillo, CA, USA). The antigen-antibody complexes were visualized using a secondary horseradish peroxidase-conjugated goat anti-mouse IgM antibody (Sigma-Aldrich) at a dilution of 1:10,000, and were detected with SuperSignal chemiluminescent substrate (Pierce Biotechnology, Rockford, IL, USA) using a Bio-Rad ChemiDox XRS. Membranes were incubated overnight at 4°C with human serum samples at a dilution of 1:200 in TBST with 5% BSA, and an anti-IgE antibody was used to detect bound antibodies.

### Deglycosylation of *Am. americanum* salivary proteins

Peptide-N-glycosidase F (PNGase F) was used for the deglycosylation of tick salivary glands and saliva glycoproteins as per the manufacturer’s instructions. Briefly, the tick protein samples (150 µg) were incubated with PNGase (15 units) at 37°C for 3 h, then the reactions were stopped by heating to 100°C for 5 min. Deglycosylation efficacy was assessed by SDS-PAGE and western blotting as described previously.

### Protein analysis

Selected bands were excised from the gels and were placed in water and shipped to MS Bioworks (Ann Arbor, MI, USA) for trypsin digestion and LC-MS/MS analysis of the resulting peptides. Gels were washed with 25 mM ammonium bicarbonate followed by acetonitrile. They were then reduced with 10 mM dithiothreitol at 60°C and were alkylated using 50 mM iodoacetamide at room temperature. Samples were then digested with trypsin (Promega, Madison, WI, USA) at 37°C for 4 h. The reaction was then quenched using formic acid. Half of each sample was analyzed using nano LC-MS/MS with the HPLC system (Waters NanoAcquity, Milford, MA, USA) interfaced to a ThermoFisher Q Exactive. Peptides were loaded on a trapping column and eluted over a 75-µm analytical column at 350 nL/min, and both columns were packed with Luna C18 resin (Phenomenex, Torrance, CA, USA). The mass spectrometer was operated in data-dependent mode, with the Orbitrap operating at 60,000 FWHM and 17,500 FWHM for MS and MS/MS, respectively. The 15 most abundant ions were selected for MS/MS. Data were searched using a local copy of Mascot against the UniProt *Ixodes scapularis* and NCBI *Amblyomma americanum* databases with monoisotopic mass values, 10 ppm peptide mass tolerance, 0.002 Da fragment mass tolerance, and maxed missed cleavages of 2. The Mascot DAT files were parsed into Scaffold (Proteome software, Portland, OR, USA) for validation, filtering, and to create a non-redundant list per sample. Data were filtered using 1% protein and peptide false discovery rate (FDR), which requires at least two unique peptides per protein.

### Immunolocalization of α-galactose

Immunolocalization studies of α-galactose were performed on partially-fed salivary glands from *Am. americanum* and *De. variabilis*. The tick salivary glands were fixed in 1× PBS containing 4% formaldehyde and were stored at 4°C. The salivary glands were permeabilized using 0.1% Triton X-100 (Sigma-Aldrich) for 30 min and then blocked in 1× PBST containing 5% bovine serum albumin (BSA) for 1 h at room temperature. Salivary glands were incubated overnight at 4°C with α-galactose IgM antibody (1:20; Enzo) in 1× PBST containing 5% BSA, after which they were incubated with an Alexa Fluor 546 goat anti-mouse IgM secondary antibody (1:100) (Life Technologies), and fluorescent dye 633-I phalloidin (1:100) (Abnova, Walnut, CA, USA) in 1× PBS containing 5% BSA for 1 h in the dark. All incubations and washes were performed on a rocking plate at room temperature unless otherwise indicated. Salivary glands were mounted on glass slides using PROLONG Gold anti-fade reagent with DAPI (Life Technologies) mounting medium. Tissues prepared in this manner were mounted and viewed under a Zeiss LSM 510 META confocal microscope running ZEN 2009 software (Zeiss, Heidelberg, Germany), using the 10×, 20×, 40×, 63×, and 100× objectives and the 405 nm, 545 nm, and 633 nm wavelength lasers.

### Magnetic pull-down of α-galactose-containing proteins

Proteins with terminal α-gal galactosylations were removed from partially-fed *Am. americanum* salivary gland tissue homogenates using DynaBeads α-mouse IgM magnetic beads (Invitrogen, Carlsbad, CA, USA). Briefly, mouse IgM-specific magnetic beads were incubated with the α-gal IgM (M86) antibody overnight on a rocker at 4°C. Antibody/bead complexes were separated from the supernatant, and tick salivary gland homogenates were added to the tube and incubated for 20 min at 4°C. The supernatant was removed and the pellet was washed three times for 5 min. Protein and antibodies were released from the beads using elution buffer. Products were run in an immunoblot assay to determine successful pull-down, and corresponding bands were excised and sent for LC-MS/MS analysis.

### N-linked glycan profiling

N-linked glycans were released from 30 µL of *Am. americanum, Am. maculatum*, and *Ix. scapularis* saliva with an estimated protein concentration of 200 µg, after being reduced, alkylated, and then digested with trypsin in Tris-HCl buffer overnight. After protease digestion, the sample was passed through a C18 sep pak cartridge, washed with 5% v/v acetic acid, and the glycopeptides were eluted with a blend of isopropanol in 5% v/v acetic acid, before being dried by SpeedVac. The dried glycopeptide eluate was treated with a combination of PNGase A (Sigma) and PNGase F (New England Biolabs, Ipswitch, MA, USA) to release the N-linked glycans. The digest was then passed through a C18 sep pak cartridge to recover the N-glycans. The N-linked glycans were then permethylated for structural characterization by mass spectrometry ^25^. Briefly, the dried eluate was dissolved with dimethyl sulfoxide and methylated with NaOH and methyl iodide. The reaction was quenched with water and per-*O*-methylated carbohydrates were extracted with methylene chloride and dried under N_2_. The permethylated glycans were reconstituted in 1:1 MeOH:H_2_O containing 1 mM NaOH, then introduced to the mass spectrometer (Thermo Fusion Tribrid Orbitrap) with direct infusion at a flow rate of 0.5 µL/min. Full MS spectra, as well as an automated “TopN” MS/MS program of the top 300 peaks, were collected and fragmented with collision-induced fragmentation. These fragmentation data were used to confirm a Hex-Hex-HexNAc signature, both with a diagnostic fragment, as well as expected neutral losses.

### Indirect basophil activation test

Peripheral blood mononuclear cells (PBMCs) taken from a healthy, non-α-gal allergic donor (α-gal sIgE < 0.10) were isolated using a Ficoll–Paque gradient (GE Healthcare, Chicago, IL, USA). Endogenous IgE was stripped from basophils within the PBMC fraction by incubating the cells with cold lactic acid buffer (13.4 mM lactic acid, 140 mM NaCl, 5mM KCl) for 15 min. Basophils were sensitized with plasma from α-gal allergic and non-allergic subjects overnight in RPMI 1640 cell culture media (Corning CellGro, Manassas, VA, USA) in the presence of IL-3 (1 ng/mL, R&D Systems, Minneapolis, MN, USA) at 37°C and 5% CO_2_.

PBMCs were subsequently stimulated for 30 min with RPMI media, cetuximab (10 μg), rabbit anti-human IgE (1 μg; Bethyl Laboratories Inc., Montgomery, TX, USA), saliva from *Am. americanum* (10 μg), or partially-fed salivary gland extracts from *Am. americanum* (50 μg), *Am. maculatum* (50 μg), or *Ix. scapularis* (50 μg). Stimulation reactions were stopped with 20 mM EDTA and PBMCs stained with fluorescently-labelled antibodies against CD123 (BioLegend, San Diego, CA, USA), human lineage 1 (CD3, CD14, CD16, CD19, CD20, CD56, BD Biosciences, San Jose, CA, USA), HLA-DR, CD63 (eBiosciences ThermoFisher, Waltham, MA, USA), and CD203c (IOTest Beckman Coulter, Marseille, France) in flow cytometry staining buffer with 0.02% NaN_3_. Samples were acquired on a CyAN ADP flow cytometer (Beckman Coulter, Brea, CA, USA) and analyzed using FlowJo v10 software (FlowJo LLC, Ashland, OR, USA). Data analysis was performed using Prism version 7.03 (GraphPad Software, La Jolla, CA, USA). Mann–Whitney U tests were used to compare the frequency of CD63+ basophils detected following stimulation with various compounds. A p-value ≤0.05 was considered significant.

## Results

### Screening of multiple tick species

The species chosen for this study were *Ix. scapularis, Am. maculatum, Am. americanum*, and *De. variabilis* because they are the most prevalent tick species in the known hotspot of AGS-reported cases. Immunoblot analysis indicated the cross-reactivity of *Ix. scapularis* unfed, partially-fed, and partially-fed *Am. americanum* salivary glands to anti-gal antibodies to high and low molecular weight tick antigens (Figure 1). Lack of α-gal cross-reactivity in the immunoblots of unfed lone-star tick tissues prompted us to determine the time-dependent presence of α-gal-containing antigens from one day post-infestation to replete tick tissues (Figure 2). Immunoblotting revealed that the expression of α-gal-containing antigens appeared in salivary tissues in a time-dependent manner throughout the feeding process. Saliva from partially-blood-fed *Am. americanum* ticks cross-reacted with anti-α-gal antibodies; however, *Am. maculatum* saliva antigens exhibited no reactivity.

**Figure 1.**
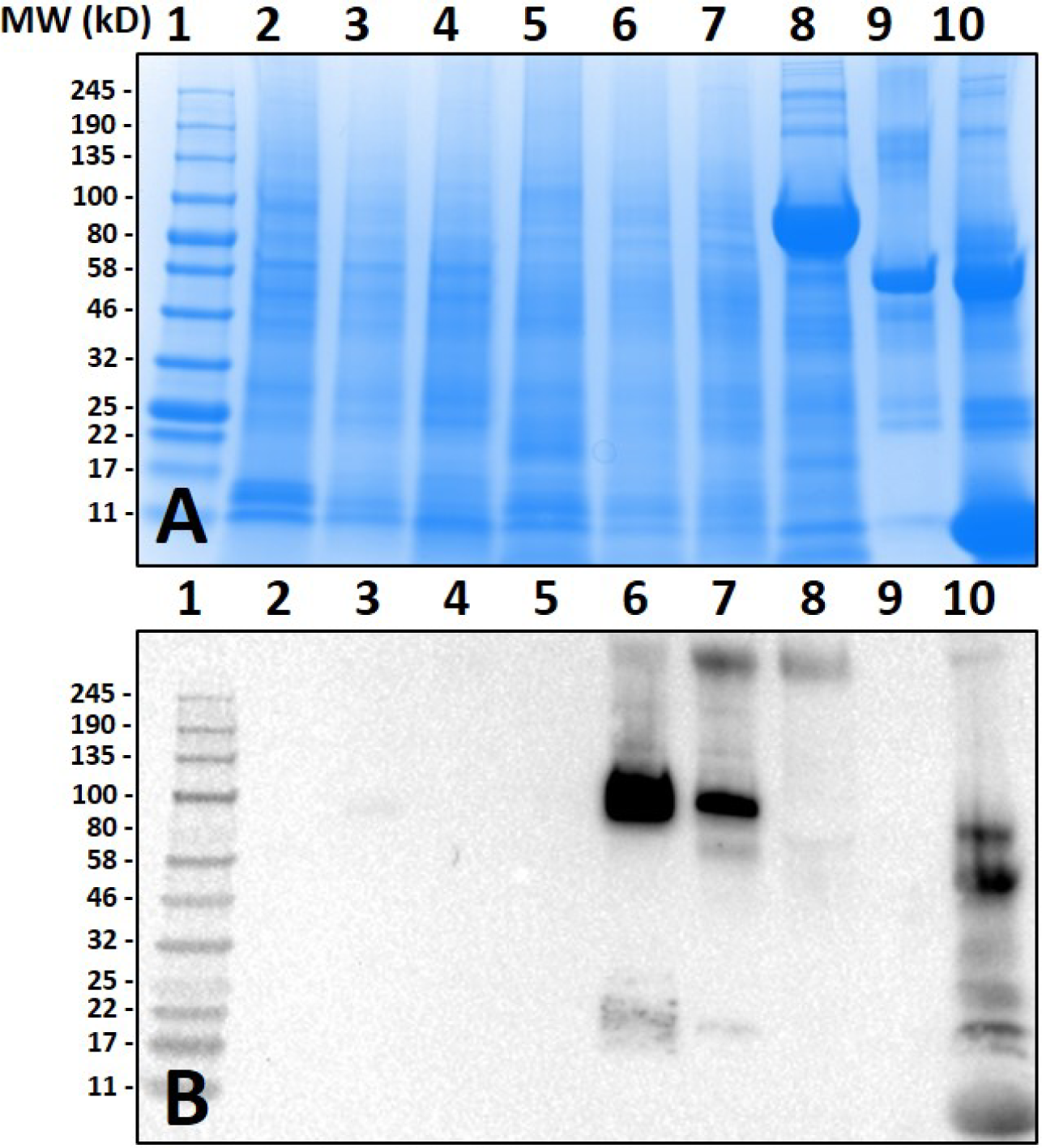
Screening of proteins from *Amblyomma americanum* (*Aam*) for □-gal. Unfed (UF) and partially-fed (PF) midgut (MG) and salivary gland (SG) tissue homogenates and saliva were fractionated on A) 7.5% SDS-PAGE, B) western blot using anti-alpha-gal antibody. Lane 1: broad range (11–245 kDa) pre-stained protein standard, Lane 2: *Aam* UF MG, Lane 3: *Aam* (3 days) PF MG, Lane 4: Aam (11 days) PF MG, Lane 5: *Aam* UF SG, Lane 6: *Aam* (3 days) PF SG, Lane 7: *Aam* (11 day) PF SG, Lane 8: *Aam* saliva, Lane 9: Bovine serum albumin and, Lane 10: Diluted sheep blood (1:100).

**Figure 2.**
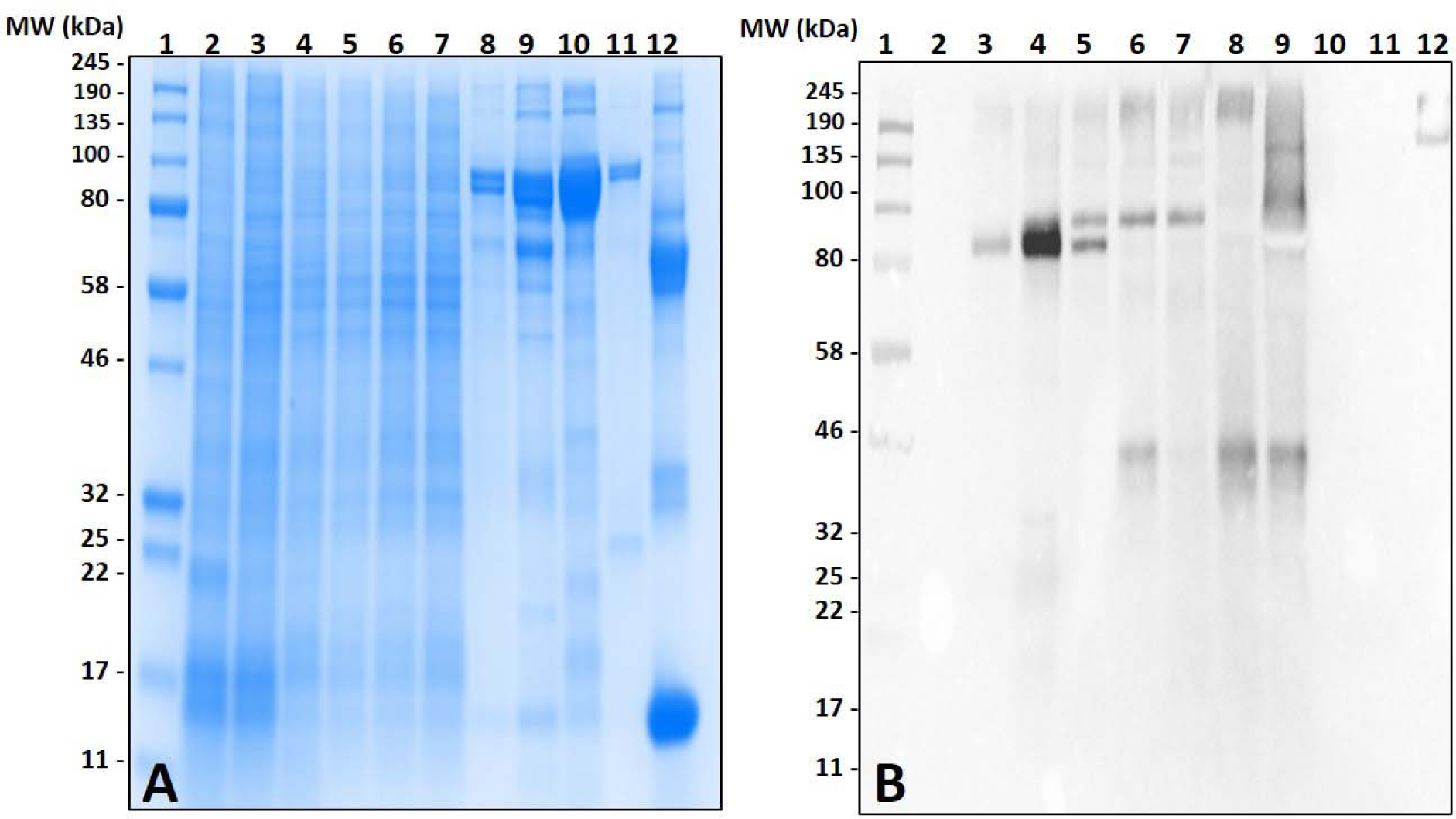
Identification of □-Gal in the salivary glands of *Amblyomma americanum* (*Aam*) during the blood meal. The unfed (UF) and partially-fed (PF) salivary glands (SG) from *Aam* throughout the blood meal were analyzed, along with saliva from *Aam* and *Amblyomma maculatum (Amac)* drooled with pilocarpine and dopamine. A) SDS-PAGE using Any kDa™ Mini-PROTEAN TGX gel, B) western blot using anti-alpha-gal antibody. Lane 1: A broad range (11–245 kDa) pre-stained protein standard, Lane 2: *Aam* UF SG, Lane 3: *Aam* (one day) fed SG, Lane 4: *Aam* (3 days) PF SG, Lane 5: *Aam* (5 days) PF SG, Lane 6: *Aam* (7 days) PF SG, Lane 7: *Aam (*8 days) PF SG, Lane 8: *Aam* saliva (Dopamine), Lane 9: *Aam* saliva (Pilocarpine), Lane 10: *Amac* saliva (Pilocarpine), Lane 11: *Amac* saliva (Dopamine) and, Lane 12: Diluted bovine blood (1:100).

Surprisingly, the cross-reactivity of *Ix. scapularis* salivary antigens differed between unfed and partially-fed tissues collected at various time points (Figure 3). These results depict the immune-reactivity of α-gal antibodies to unfed tissue antigen sizes ranging from 32–50 kDa, and 245 kDa and higher molecular weights. However, blood meal induces salivary antigens to cross-react with α-gal antibodies consistently in the range of 100–135 kDa (Figure 3B). Interestingly, α-gal antibodies cross-reacted with unfed midgut tissue antigens of *Ix. scapularis*, and upon blood feeding this reactivity disappeared (Figure 3D).

**Figure 3.**
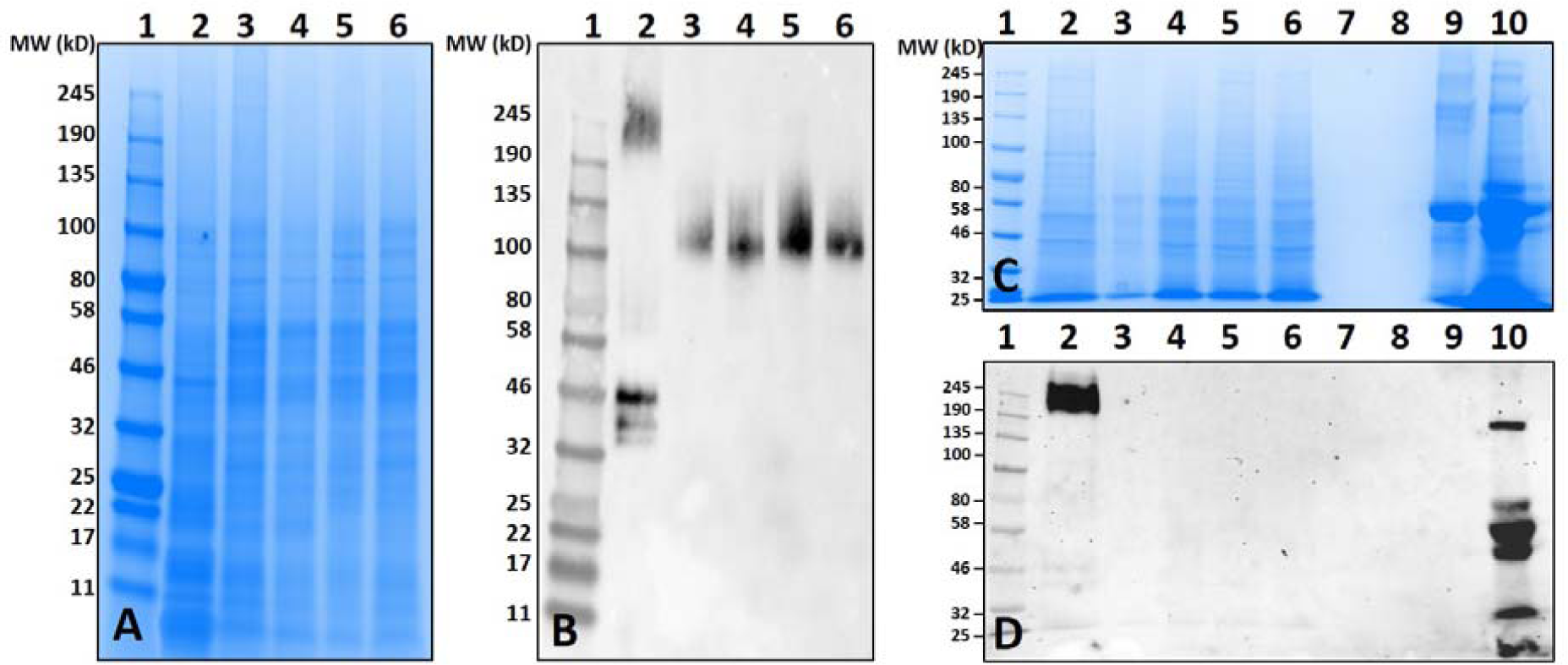
Identification of □-Gal in the salivary gland (SG) and midgut (MG) tissues of *Ixodes scapularis* (*Ixscap*) during the blood meal. A) SDS-PAGE of *Ixscap* SG using 7.5% Mini-PROTEAN TGX, B) western blot using anti-alpha-gal antibody. Lane 1: A broad range (11–245 kDa) prestained protein standard, Lane 2: *Ixscap* UF SG, Lane 3: *Ixscap* (3 days) PF SG, Lane 4: *Ixscap* (5 days) PF SG, Lane 5: *Ixscap* (6 Days) PF SG, Lane 6: *Ixscap (*8 days) PF SG. C) SDS-PAGE of *Ixscap* MG tissues. Lane 1: A broad range (11–245 kDa) prestained standard, Lane 2: *Ixscap* UF MG, Lane 3: *Ixscap* (3 days) PF MG, Lane 4: *Ixscap (*5 days) PF MG, Lane 5: *Ixscap* (6 days) PF MG, Lane 6: *Ixscap (*8 days) PF MG, Lane 7-8: blank. Lane 9: Bovine serum albumin and, Lane 10: Diluted sheep blood (1:100). D) Western blot using anti-alpha-gal antibody.

Intriguingly, unfed *Am. americanum* salivary glands did not show any cross-reactivity with anti-α-gal antibodies. Both unfed and partially-blood fed *De. variabilis* and *Am. maculatum* salivary glands also lacked immune-reactivity with α-gal antibodies. The western blot was normalized using an antibody against beta-actin, and the cross-reacting bands containing α-gal-linked antigens were excised and peptides identified using LC-MS/MS. These data suggest that some tick species may contain α-gal-like glycans in their salivary gland tissues. Gut tissues of tested tick species showed no cross-reactivity with the test antibodies.

Mass spectrometry (LC/MS-MS) analysis of gel excisions (Figure 4A) revealed numerous protein peptides of various functions (Table 1 and Supplementary Data Table S1). The excision of *Am. americanum* salivary glands contained peptides from proteins such as an abundant hemelipoprotein precursor that contains a VWD domain, glucose-regulated protein grp-94/endoplasmin hsp90 family, and endoplasmic reticulum resident protein glycosyltransferases. Unfed *Ix. scapularis* contained multiple glycoside hydrolases including alpha-L-fucosidase and alpha-D-galactosidase, enzymes known to cleave terminal alpha-L-fucosides and alpha-D-galactosides. The unfed salivary glands also contained numerous lectins including galectin, hemolectin, and a mannose-binding endoplasmic reticulum-Golgi compartment lectin. In the unfed *Ix. scapularis* salivary glands, we also observed oligosaccharyl transferases, a sugar transporter protein capable of transporting galactose, heat shock proteins, hemomucin, heme lipoproteins or heme lipoglycoproteins, and ixoderin B. In the partially-fed *Ix. scapularis* salivary glands, we observed glycoside hydrolases including alpha-mannosidases, alpha-glucosidases, lysosomal glucosidase, and glucosidase II containing a galactose mutarotase domain. Additionally, there were a few glycosyltransferases including oligosaccharyltransferase and alpha-1,3-glucosyltransferase.

**Figure 4.**
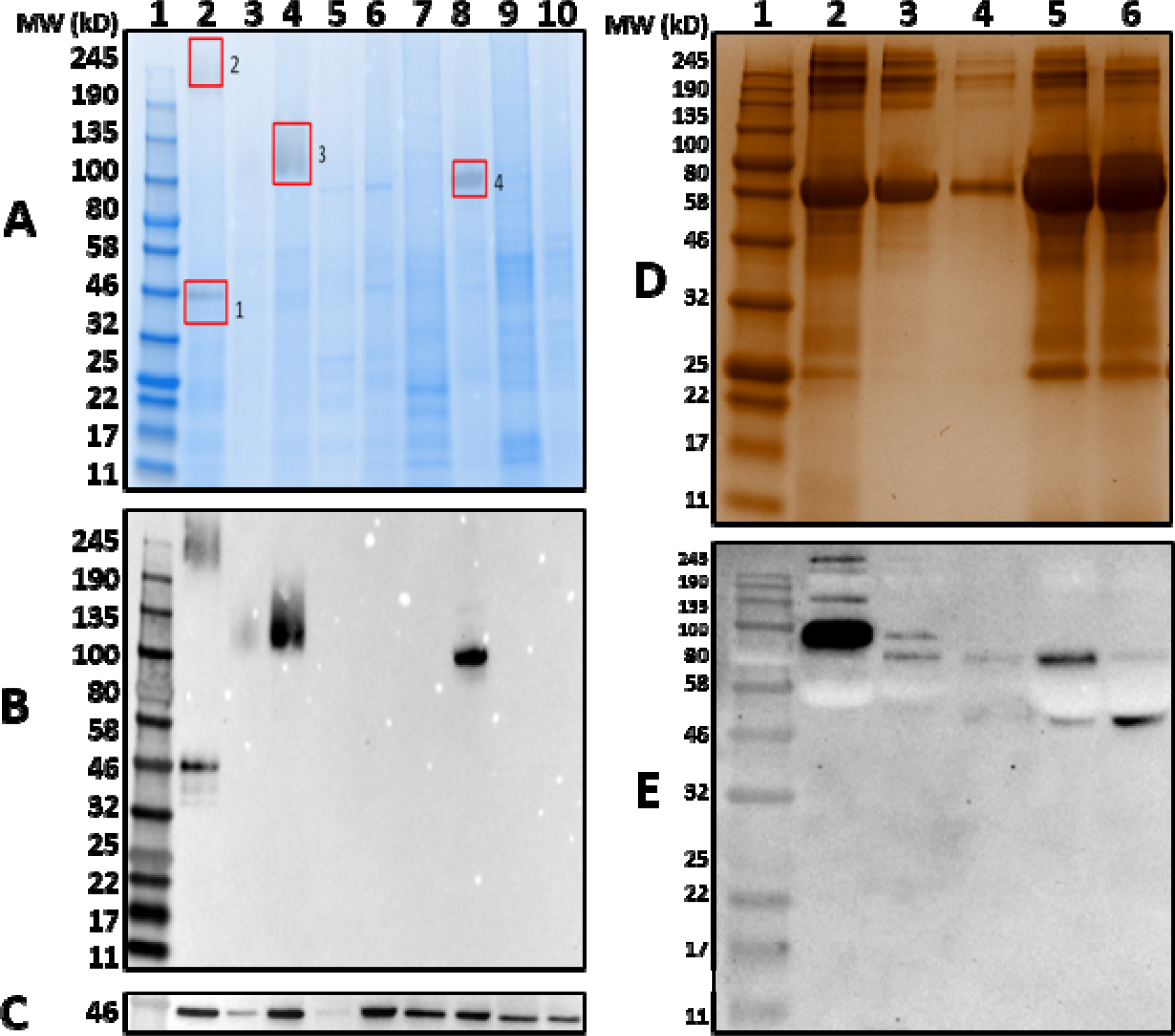
Identification of α-gal-containing salivary antigens from various tick species using immunoproteome workflow. A total of 15 *µ*g of Unfed (UF) and partially-fed (PF) salivary gland (SG) tissue homogenates from *Ix. scapularis (Ixscap), Am. maculatum (Amac), Am. americanum (Aam)*, and *De. variabilis (Dvari)* were fractionated using A) an overlay of the SDS-gel and blot showing excised bands for mass spectrometry analysis. Lane 1: broad range pre-stained protein standard. Lane 2: *Ixscap* UF SG. Lane 3: overflow. Lane 4: *Ixscap* PF SG. Lane 5: *Amac* UF SG. Lane 6: *Amac* PF SG. Lane 7: *Aam* UF SG. Lane 8: *Aam* PF SG. Lane 9: *Dvari* UF SG. Lane 10: *Dvari* PF SG. B) Western blot using anti-alpha-gal antibody, C) western blot probed using Beta-actin monoclonal antibody, D) silver-stained gel of magnetic pull-down of α-gal-containing proteins using magnetic beads. Lane 1: broad range molecular weight protein standard, lane 2: “depleted” *Aam* SG, lane 3: the α-gal pull-down product, lane 4: the magnetic beads after incubation with *Aam* SG, lane 5: “depleted” α-gal-specific IgM, and lane 6: α-gal-specific IgM. E) Western blotting using anti-gal antibody.

**Table 1.**
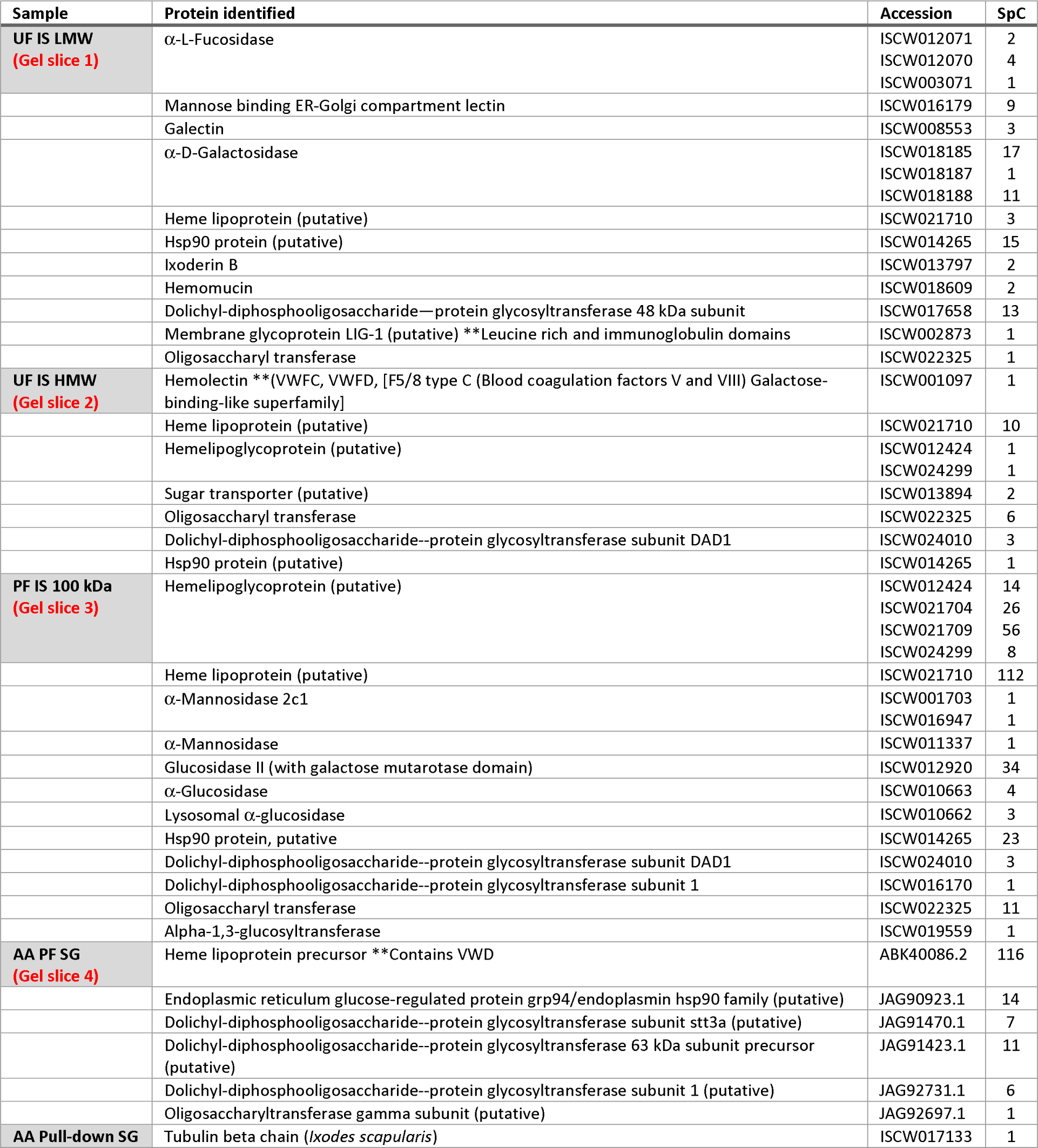
Select proteins identified from mass spectrometry of gel slices and magnetic pull-down assay. Samples, protein names, accession numbers, and exclusive spectrum counts (SpC) are listed below (full list of proteins included in spreadsheet).

### Magnetic pull-down and the proteomics of tick salivary glands

Previous proteomic experiments using excised bands provided insight into the tick antigens containing α-gal in salivary gland tissues but narrowing this down to unknown proteins based on surface glycosylations presents its own technical challenges and difficulties. IgM-specific magnetic beads were used to capture antigens with alpha-galactosyl residues using the anti-gal antibody. The quantities of captured antigens were expected to be low, and therefore silver staining was used to visualize antigens (Figure 4D). To further verify the specific binding of the captured antigens in the “pull-down” and “depleted” fractions, α-gal antibody was used to confirm the cross-reactivity (Figure 4E). The “depleted” fraction showed the usual cross-reactivity at ∼95 kDa, and the “pull-down” fraction showed a band with less intensity at a similar molecular weight. It is likely that the bands at ∼80 kDa and ∼50–55 kDa are artifacts of the IgM antibodies that were not removed from the α-galactosyl-containing proteins. The magnetic pull-down assay was able to capture the α-gal-containing epitope from *Am. americanum* salivary gland tissue homogenates and the product was excised and LC-MS/MS analysis identified the tubulin β-chain of *Am. americanum* (A0A0C9SCB7).

### Artificial feeding of ticks and deglycosylation of salivary proteins

Non-primate mammals contain an abundance of α-gal, and tick tissues used in these experiments were from ticks that were fed on mammals. Therefore, it is possible that the detected α-gal-containing antigens from post-attachment ticks could have originated from those animals, even though the previous data indicated that only two species of tick, *Ix. scapularis* and *Am. americanum*, had detectable quantities of α-gal post-feeding. To determine whether the source of α-gal is host blood or if α-gal is being synthesized by the tick, *Am. americanum* females were fed *in vitro* using an artificial membrane feeding system that contained defibrinated human blood. Western blot analysis showed the cross-reactivity of α-gal-containing antigens fed on both sheep and human blood (Figure 5). Noticeably, tick antigens ranging from 80–100 kDa in size from the salivary glands fed on human blood cross-reacted with α-gal antibodies. The results from this experiment support the argument that α-gal is being synthesized by the tick via a so far unknown mechanism.

**Figure 5.**
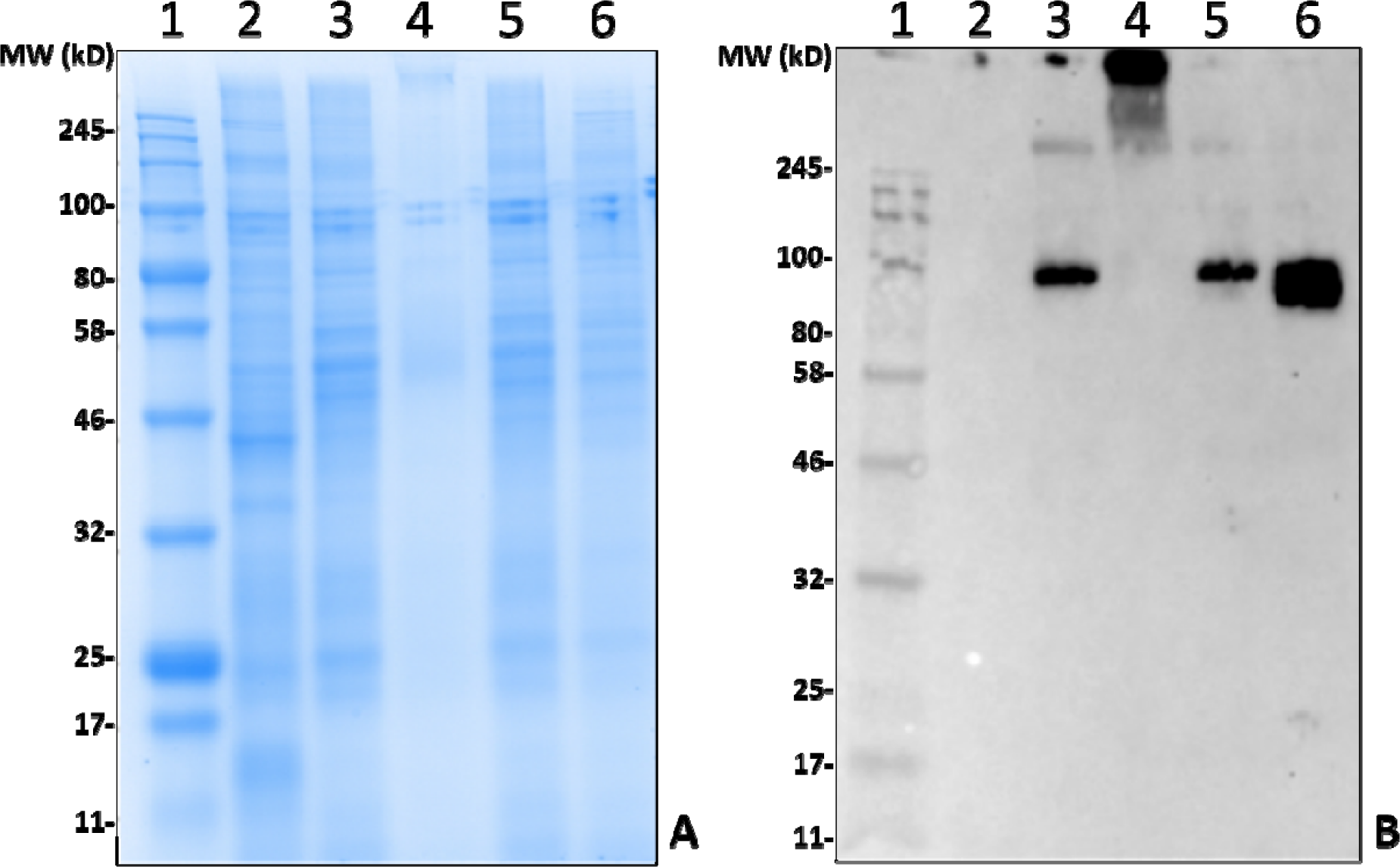
Deglycosylated and human blood-fed *Amblyomma americanum* (*Aam*) salivary gland (SG) analysis. SDS-PAGE (A) and *α*-gal immunoblot (B). Lane 1: a broad range molecular weight ladder, Lane 2: *Aam* UF SG. Lane 3: 7-*Aam* (7 days) PF SG. Lane 4: deglycosylated *Aam* (7 days) PF SG. Lane 5: *Aam* (7 days) PF SG. Lane 6: Aam SG that were fed defibrinated whole human blood using an artificial membrane feeding system.

While our data revealed that anti-gal antibodies are cross-reacting with α-galactosyl epitopes on tick salivary antigens, the identity of the glycosylated protein or lipid of interest remains uncertain. To narrow down the possibility of this being part of a bacterial lipopolysaccharide or a lipid-linked glycan, *Am. americanum* salivary gland protein isolates were incubated in PNGase F to cleave the N-linked glycan chain at asparagine residues (Figure 5). Salivary glands treated with PNGase F were compared with untreated salivary glands, and the anti-gal antibody was visualized above the uppermost limit of the gel/membrane, which indicates that there was successful cleavage of the glycans. Together, these data suggest that the anti-gal antibody is binding to N-linked glycans found in the partially-fed *Am. americanum* salivary glands.

### Immunolocalization of α-gal in tick salivary gland tissues

Immunolocalization of α-gal-containing antigens was conducted using partially-blood-fed salivary glands of *Am. americanum, Ix. scapularis,* and *Am. maculatum* to determine the subcellular location of the α-gal moiety in tick tissues. The results indicated the presence of detectable α-gal-containing antigens in the salivary gland acini in proximity to the secretory vesicle in partially-blood-fed *Am. americanum* and *Ix. scapularis* (Figure 6). As expected, no cross-reactivity of anti-gal antibody was noted in *Am. maculatum*, which corresponds with the results of immunoblot analysis (Figure 4 and Supplement 1).

**Figure 6.**
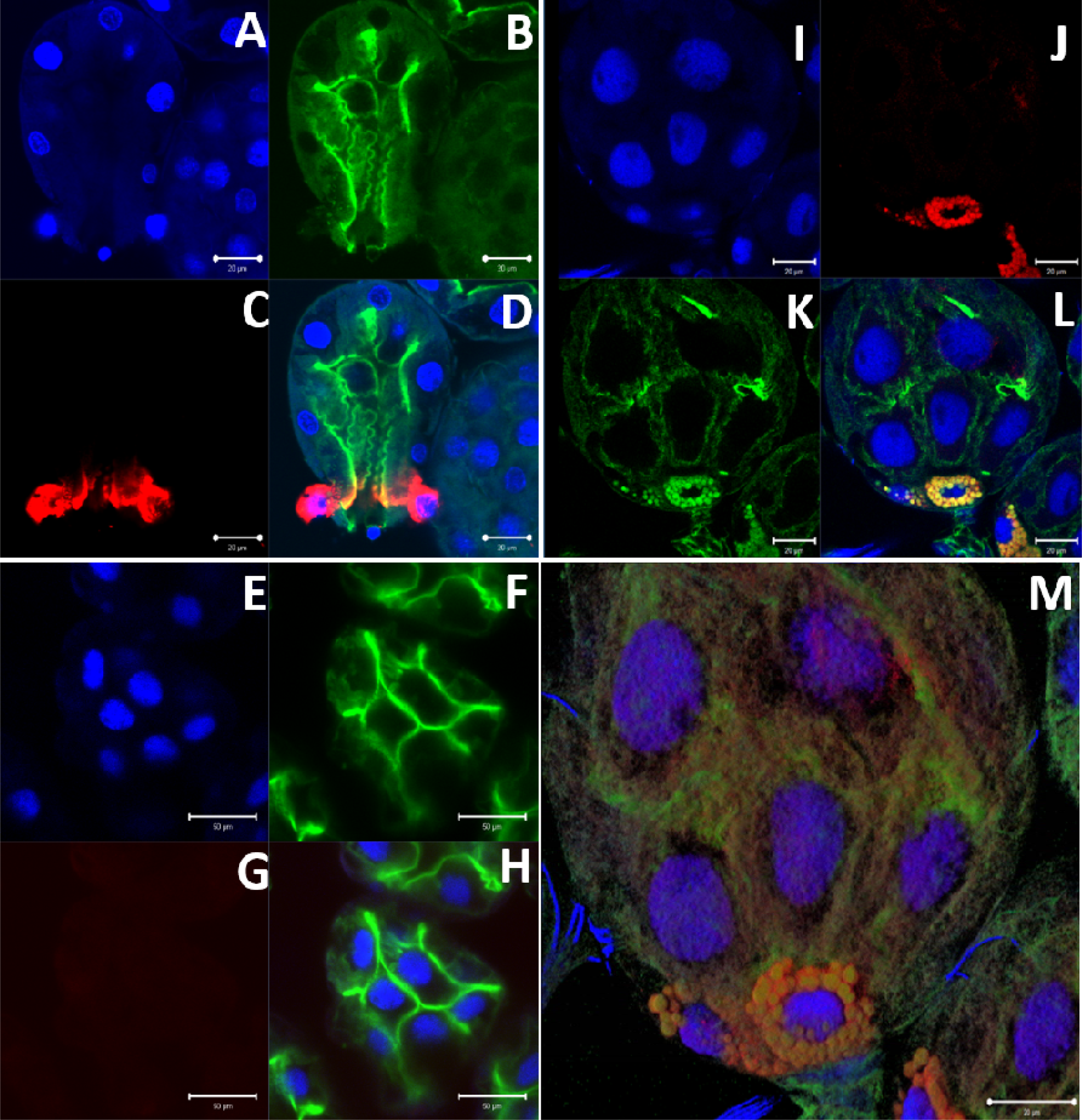
α-Gal immunolocalization in partially-fed *Ix. scapularis* (*Ixscap*), *Am. maculatum* (*Amac*), and *Am. americanum* (*Aam*) salivary glands (SG). *Ixscap* SG images (63×) using DAPI (BLUE Color) (A), phalloidin-stained F-actin (Green Color) (B), alpha-gal antibody (Red Color) (C), and merged images (D). *Amac* SG images (40×) using DAPI (E), phalloidin-stained F-actin (F), alpha-gal antibody (G), and merged images (H). *Aam* SG images (100×) using DAPI (I), alpha-gal antibody (J), phalloidin-stained F-actin (K), merged images (L), and 100× 3D projection from Z-stack (M).

### N-linked glycan profile of tick saliva and salivary gland tissues

N-linked glycan profiling of *Am. maculatum, Am. americanum,* and *Ix. scapularis* revealed two critical insights into the glycosylation of tick salivary proteins (Table 2 and 3). First, the unfed salivary glands of both *Am. americanum* and *Am. maculatum* contained no detectable amounts of α-gal, but glycans in the unfed salivary glands of *Ix. scapularis* contained α-gal. Secondly, the partially-fed salivary glands and saliva from *Am. maculatum* showed no detectable α-gal, which corresponds to our immunoblot and immunolocalization experiments; however, *Am. americanum* and *Ix. scapularis* contained detectable quantities and multiple glycoforms of α-gal (Table 2). The overall abundance of α-gal glycoforms in *Am. americanum* partially-fed salivary glands tested was greater than 1.12% of the total N-glycans detected, but the saliva consisted of approximately 0.15% of the total N-glycans detected. *Ix. scapularis* unfed salivary gland N-glycans containing α-gal comprised 6.3% of overall N-glycans detected, and more than 1.7% in the partially-fed salivary glands, but only trace amounts, below the quantifiable limit, of α-gal glycans were found within the saliva. The majority of the glycoforms identified were biantennary extended-galactose structures that had core fucosylations, and a few were identified as single-antennary species (hybrid-type) in the non-fucosylated form. This information correlates with the results of immunoblotting experiments and strengthens the theory that some tick species can acquire or synthesize α-gal, while others lack this capacity. Comprehensive details of all identified N-linked glycoforms have been provided (Supplementary Data Table S2).

**Table 2.**
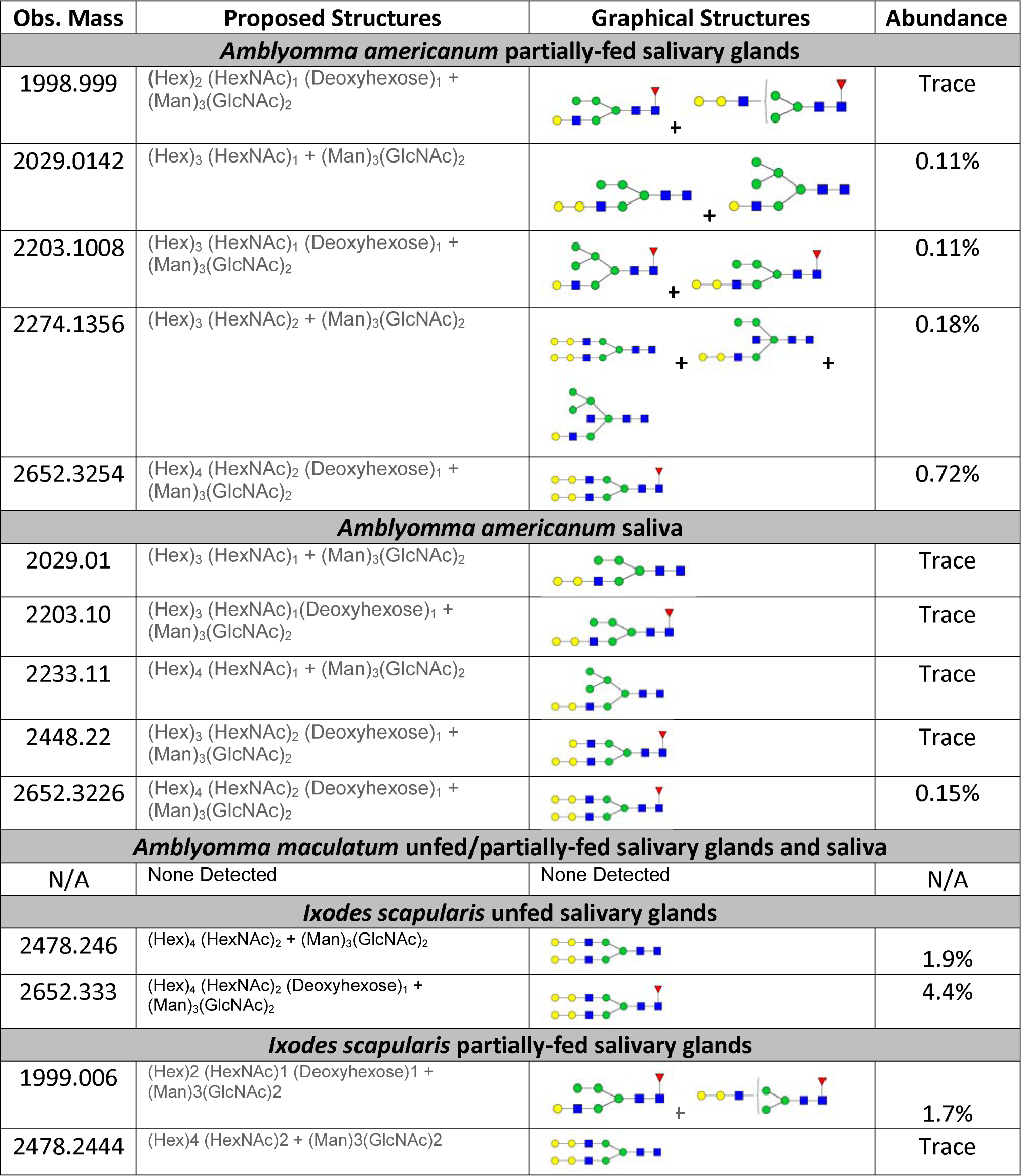
Quantification of alpha-galactosyl-containing N-linked glycans by NSI-FTMS. Glycoforms with alpha-galactosyl epitope observed in partially-fed *Amblyomma americanum* salivary glands and saliva, and *Ixodes scapularis* unfed and partially-fed salivary glands.

| Obs. Mass | Proposed Structures | Graphical Structures | Abundance |
| --- | --- | --- | --- |
| ***Amblyomma americanum* partially-fed salivary glands** |  |  |  |
| 1998.999 | (Hex)<sub>2</sub> (HexNAc)<sub>1</sub> (Deoxyhexose)<sub>1</sub> + (Man)<sub>3</sub>(GlcNAc)<sub>2</sub> |  | Trace |
| 2029.0142 | (Hex)<sub>3</sub> (HexNAc)<sub>1</sub> + (Man)<sub>3</sub>(GlcNAc)<sub>2</sub> |  | 0.11% |
| 2203.1008 | (Hex)<sub>3</sub> (HexNAc)<sub>1</sub> (Deoxyhexose)<sub>1</sub> + (Man)<sub>3</sub>(GlcNAc)<sub>2</sub> |  | 0.11% |
| 2274.1356 | (Hex)<sub>3</sub> (HexNAc)<sub>2</sub> + (Man)<sub>3</sub>(GlcNAc)<sub>2</sub> |  | 0.18% |
| 2652.3254 | (Hex)<sub>4</sub> (HexNAc)<sub>2</sub> (Deoxyhexose)<sub>1</sub> + (Man)<sub>3</sub>(GlcNAc)<sub>2</sub> |  | 0.72% |
| ***Amblyomma americanum* saliva** |  |  |  |
| 2029.01 | (Hex)<sub>3</sub> (HexNAc)<sub>1</sub> + (Man)<sub>3</sub>(GlcNAc)<sub>2</sub> |  | Trace |
| 2203.10 | (Hex)<sub>3</sub> (HexNAc)<sub>1</sub>(Deoxyhexose)<sub>1</sub> + (Man)<sub>3</sub>(GlcNAc)<sub>2</sub> |  | Trace |
| 2233.11 | (Hex)<sub>4</sub> (HexNAc)<sub>1</sub> + (Man)<sub>3</sub>(GlcNAc)<sub>2</sub> |  | Trace |
| 2448.22 | (Hex)<sub>3</sub> (HexNAc)<sub>2</sub> (Deoxyhexose)<sub>1</sub> + (Man)<sub>3</sub>(GlcNAc)<sub>2</sub> |  | Trace |
| 2652.3226 | (Hex)<sub>4</sub> (HexNAc)<sub>2</sub> (Deoxyhexose)<sub>1</sub> + (Man)<sub>3</sub>(GlcNAc)<sub>2</sub> |  | 0.15% |
| ***Amblyomma maculatum* unfed/partially-fed salivary glands and saliva** |  |  |  |
| N/A | None Detected | None Detected | N/A |
| ***Ixodes scapularis* unfed salivary glands** |  |  |  |
| 2478.246 | (Hex)<sub>4</sub> (HexNAc)<sub>2</sub> + (Man)<sub>3</sub>(GlcNAc)<sub>2</sub> |  | 1.9% |
| 2652.333 | (Hex)<sub>4</sub> (HexNAc)<sub>2</sub> (Deoxyhexose)<sub>1</sub> + (Man)<sub>3</sub>(GlcNAc)<sub>2</sub> |  | 4.4% |
| ***Ixodes scapularis* partially-fed salivary glands** |  |  |  |
| 1999.006 | (Hex)<sub>2</sub> (HexNAc)<sub>1</sub> (Deoxyhexose)<sub>1</sub> + (Man)<sub>3</sub>(GlcNAc)<sub>2</sub> |  | 1.7% |
| 2478.2444 | (Hex)<sub>4</sub> (HexNAc)<sub>2</sub> + (Man)<sub>3</sub>(GlcNAc)<sub>2</sub> |  | Trace |

**Table 3.**
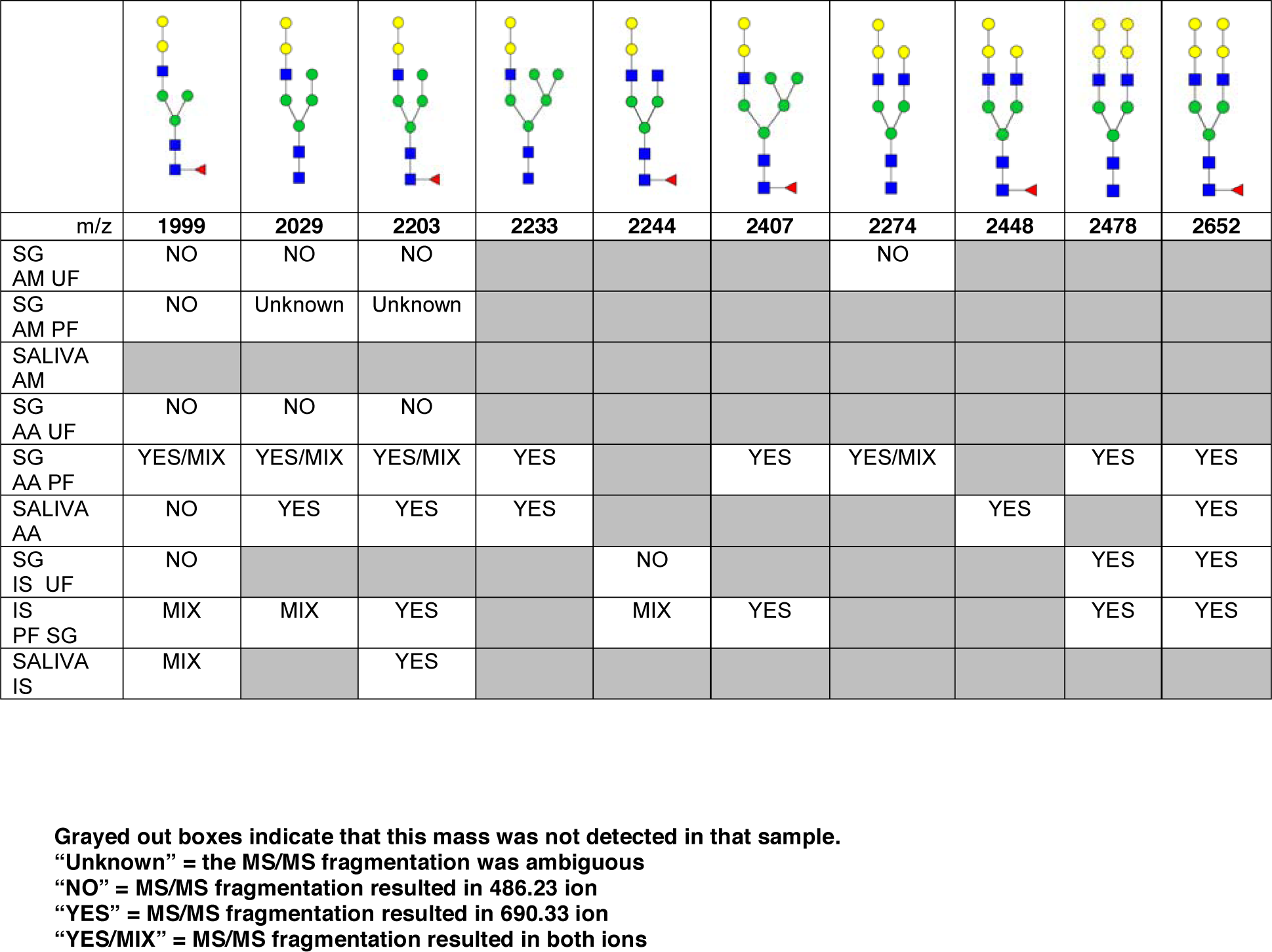
Summary of potential α-gal-containing glycoforms determined by MS/MS analysis in *Am. maculatum, Am. americanum*, and *Ix. scapularis* salivary glands and saliva.

In addition to glycoforms containing α-gal in these samples, multiple pentose-containing species were observed in all three species and in all feeding states, in both salivary glands as well as saliva. While these results do not reflect compositional analysis, MS/MS fragmentation revealed that the pentose was core-mannose attached, similar to xylose-containing structures frequently found in plants. These observed glycoforms were primarily complex-type.

### Basophil activation with tick salivary samples

Because glycan profiling demonstrated the presence of α-gal in salivary samples from *Am. americanum* and *Ix. scapularis* but not *Am. maculatum*, we sought to determine 1) if salivary α-gal moieties could activate basophils primed with α-gal sIgE, and 2) whether activation might reflect species-specific differences in α-gal glycan content. Donor basophils from a healthy, non-allergic control were stripped of IgE and primed overnight with plasma from a subject with α-gal syndrome (α-gal sIgE = 31.3 IU/mL, total IgE = 233 IU/mL). Sensitized cells were exposed to one of the following stimuli for 30 min: RPMI media, crosslinking anti-IgE antibody (positive control), α-gal-containing glycoprotein cetuximab (α-gal positive control), *Am. americanum* saliva, *Am. americanum* partially-fed salivary gland (PF SG) extract, *Ix. scapularis* PF SG extract, or *Am. maculatum* PF SG extract. CD63 expression on lineage-HLA-DR-CD123+CD203c+ basophils was assessed by flow cytometry (Figure 7). We found that the frequency of CD63+ basophils was significantly increased following sensitization with α-gal allergic plasma and stimulation with α-gal-containing tick salivary samples from *Am. americanum* (saliva and PF SG extract) and *Ix. scapularis* (p<0.05 vs. media, Figure 7). Alternatively, salivary samples from *Am. maculatum* caused small but non-significant increases in CD63+ basophils when the results of all experiments (n=3) were included. Stimulation with PF SG extract from *Ix. scapularis* produced the largest increase in CD63+ basophils, which was consistent with the high level of α-gal content detected via glycan analysis.

**Figure 7.**
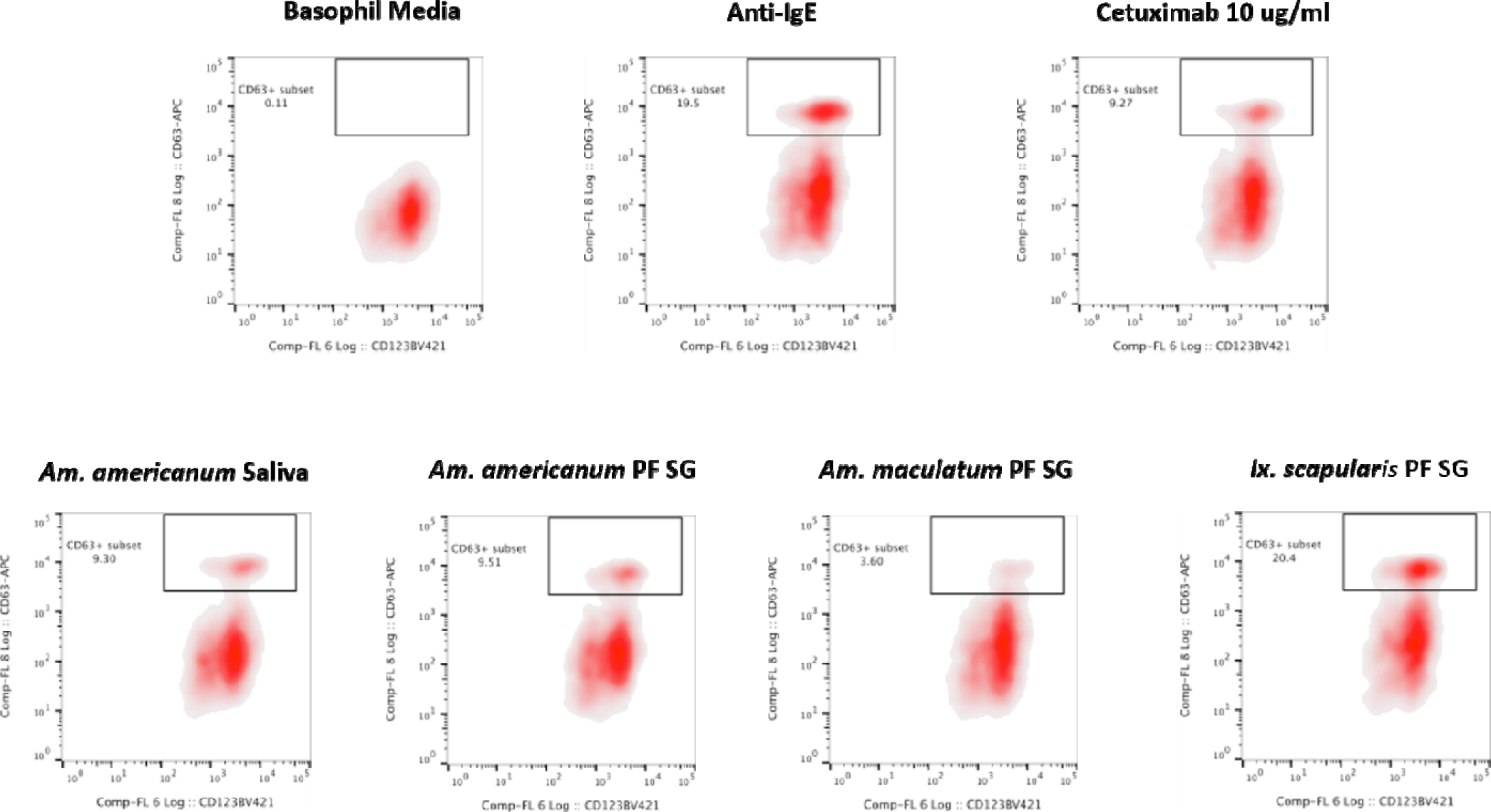
Flow cytometric analysis of human basophil activation by tick salivary proteins. Donor basophils from a healthy, non-allergic control were stripped of IgE and primed overnight with plasma from a subject with α-gal syndrome (α-gal sIgE = 31.3 IU/mL, total IgE = 233 IU/mL). Sensitized cells were exposed to one of the following stimuli for 30 min: RPMI media, crosslinking anti-IgE antibody (positive control), α-gal-containing glycoprotein cetuximab (α-gal positive control), *Am. americanum* saliva, *Am. americanum* partially-fed salivary gland (PF SG) extract, *Ix. scapularis* PF SG extract, or *Am. maculatum* PF SG extract. CD63 expression on lineage-HLA-DR-CD123+CD203c+ basophils was assessed by flow cytometry.

## Discussion

The lone-star tick has expanded its geographic range from Southwest to the East Coast of the United States. It is a vector for diseases such as spotted fever group rickettsiosis, human monocytic ehrlichiosis, southern-tick-associated rash illness, tularemia, Heartland virus infection, and infection with newly discovered Tacaribe virus [^26^–^30^]. In addition to these diseases, this tick species has been associated with delayed anaphylaxis to red meat and is the first example of a blood-feeding ectoparasite causing food allergy in the United States^5,31^. In fact, a growing body of literature suggests that bites from the lone-star tick (*Am. americanum*) are causing α-gal syndrome^5,7,8,32^. It remained unknown whether bites from *Am. americanum* trigger the development of α- gal sIgE in humans due to the presence of α-gal moieties in tick saliva or IgE arises as a class-switched anti-gal response after ecto-parasitic feeding. In this study, we identified the presence of α-gal in *Am. americanum* and the black-legged tick, *Ix. scapularis.* Furthermore, a previous study in Europe used immunohistochemical staining to show the cross-reactivity of the α-galactosyl epitope in the gut tissues of *Ix. ricinus*^11^. Our study focused on unfed and partially-fed salivary glands because they produce, contain, and secrete saliva that can be injected directly into the host^33^. In theory, if there is an absence of α-galactosyl epitopes in tick salivary glands, which secrete saliva during prolonged feeding, it is highly unlikely that α-gal will be found in the tick saliva and, therefore, α-gal sIgE would most likely reflect a Th2-driven class-switch of the ongoing anti-gal response present in all immunocompetent humans.

However, as our immunoblotting results showed, the saliva and salivary glands of *Am. americanum* female ticks express α-gal-containing antigens in a time-dependent manner throughout prolonged blood feeding (Figure 1 and 2). The presence of α-gal-containing antigens in unfed and partially-blood-fed *Ix. scapularis* was evident (Figure 3 and 4). However, the Gulf-Coast tick, *Am. maculatum*, and the American dog tick, *De. variabilis*, lacked the presence of α-gal-containing antigens (Figure 4 and Supplement 1). Unlike *Ix. scapularis* unfed midgut tissues, *Am. maculatum* and *Am. americanum* tick species showed no cross-reactivity with α-gal antibodies (Figure 1 and Supplement 1). The presence of α-gal cross-reactivity with unfed *Ix. scapularis* was not surprising as it has been reported in sister tick species *Ix. ricinus* using immunohistochemical techniques^11^ and in N-glycan profiling studies^34^. Additionally, the results of basophil activation showed high levels of CD63+ expression following stimulation using *Ix. scapularis* salivary extracts. The initial presence of α-gal-containing antigens in the unfed *Ix. scapularis* midgut and salivary glands is possibly a remnant from a blood meal during the previous life-stage before molting to the adult stage, or the ticks might have cleaved and incorporated the glycans into their own proteins. Interestingly, unfed and partially-blood-fed *Am. americanum* males showed a lack of cross-reactivity with α-gal antibodies (our unpublished data). Together, these results lead us to the conclusion that female *Am. americanum* and *Ix. scapularis* ticks express α-gal-containing proteins and might possibly use α-galactose to facilitate successful hematophagy. While it might be possible for the tick to sequester α-gal or the enzymes required to synthesize α-gal from the host during the immature developmental stages (larval or nymphal ticks), adult unfed *Am. americanum* females do not have detectable quantities of α-gal validated by immunoblotting and MALDI-TOF/TOF-MS. However, *Ix. scapularis* unfed females do have detectable quantities of α-gal in the salivary gland tissues, possibly remaining from a previous blood meal. The size of α-gal-containing antigens differs between unfed *Ix. scapularis* salivary glands and partially-blood-fed salivary glands. The switching of α- gal-containing antigens from unfed to fed salivary glands might be a strategy to remain successfully attached to the host for a prolonged period of time. Figure 8 is a reference diagram that shows which methods were used to validate the presence of α-gal in tick tissues and saliva.

**Figure 8.**
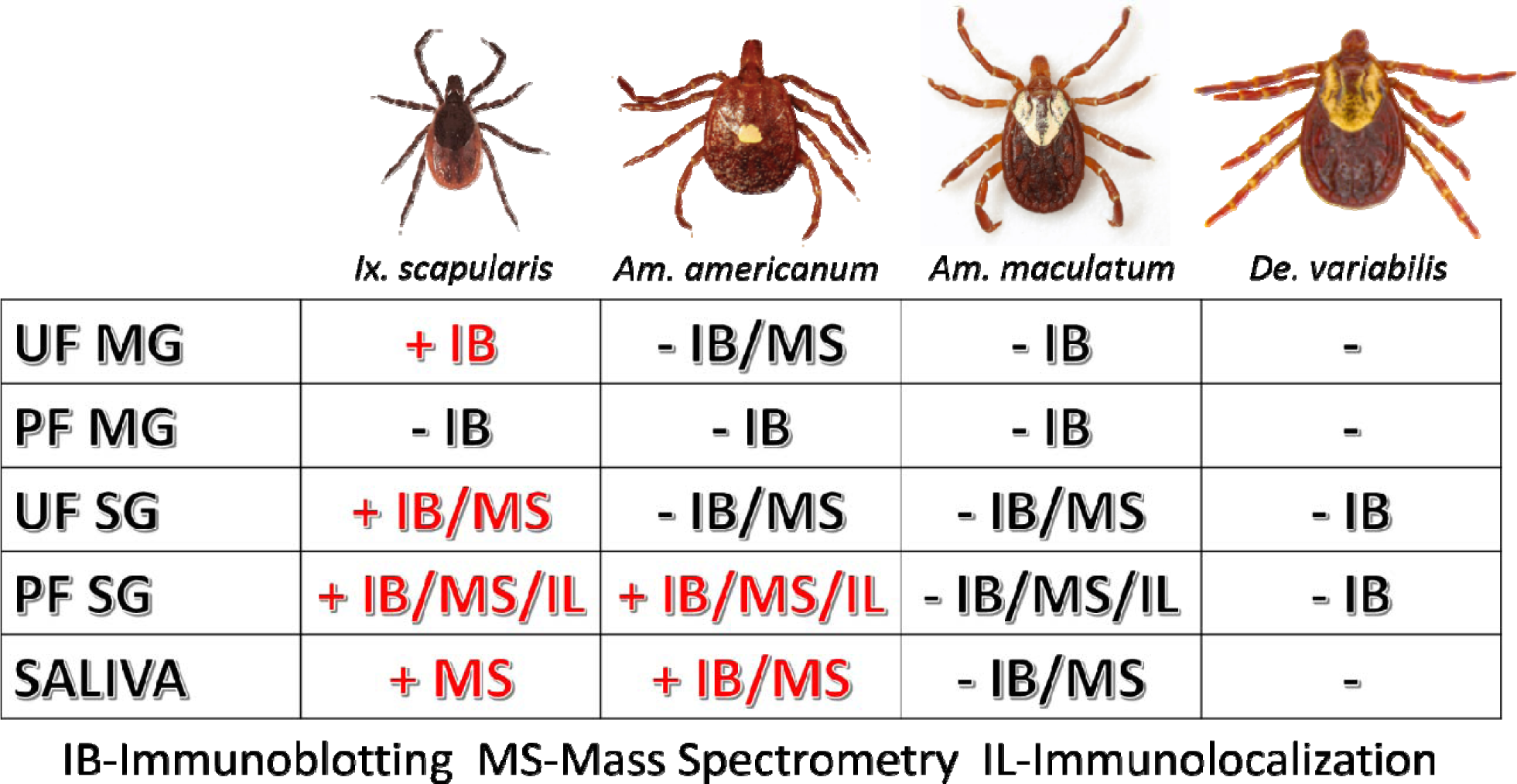
Table of tissues testing positive for α-gal in this study. The above figure shows which methods validated the presence of α-gal in individual tick tissues at various feeding stages and in the saliva. Samples that contained α-gal are denoted with “+” and red lettering, and tissues and saliva lacking α-gal are denoted with “-” and black lettering.

Incubation of *Am. americanum* salivary gland tissues with PNGase F cleaves internal glycoside bonds of asparagine-linked oligosaccharides. The results indicated that N-linked glycans with terminal α-gal caps were removed from tick salivary proteins (Figure as α-gal moieties were detected with the M86 antibody above the limit of the protein standard in the polyacrylamide gel that failed to migrate after treatment with PNGase F, as they are uncharged carbohydrate chains. This is important because it reveals that the detected α-gal is part of an N-linked oligosaccharide linked to a glycoprotein within the tick salivary glands, as opposed to an O-linked structure such as a lipopolysaccharide.

Our experiments analyzed tissues and saliva that were acquired from ticks fed on mammals that produce α-gal, and thus the host could have served as a potential source of the α-gal carbohydrates detected. To determine if the mammalian host was the source of α-gal carbohydrates, ticks were artificially fed with a membrane feeding system using human blood as a meal source, which is known to be free of the enzyme N-acetyllactosaminide alpha-1,3-galactosyltransferase^15^ and α-gal. However, *Am. americanum* ticks that were artificially fed with human blood still contained α-gal at the same molecular weight as ticks fed sheep blood, as determined using the α-gal M86 IgM antibody (Figure 5B). These results indicate that ticks possibly recycle host glycans or synthesize α-gal using alternative methods. Because there is no evidence that ticks have the necessary galactosyltransferase to produce α-gal, it is possible that the ticks use a fucosidase to cleave fucose residues from human type B blood to produce the α- gal antigens, which could then be cleaved and incorporated into tick salivary proteins or directly secreted back to the host. Another potential source of α-gal could come from bacterial galactosyltransferase enzymes that are used during cell wall biosynthesis^35^. Combined, these results suggest that ticks do not need to feed on a lower (non-primate) mammal to introduce salivary glycoproteins containing α-gal into humans.

To localize α-gal in tick salivary glands, confocal fluorescence microscopy was ustilized to visualize the emission of secondary antibodies against α-gal IgM. We primarily focused on the partially-fed stage of *Am. americanum* salivary glands, but also screened *Ix. scapularis* and *Am. maculatum* for α-gal. The images provided evidence that terminal α-gal residues on salivary glycoproteins are not found in all ticks, as α-gal was present in *Am. americanum* (Figure 1 and 2) and *Ix. scapularis* (Figure 3), but absent in *Am. maculatum* (Supplement 1). In *Am. americanum* tissues, immunolocalization of α-gal residues was primarily observed with secretory vesicles from ticks that were in the partially-fed state (Figure 6). The presence of terminal α-gal residues near secretory vesicles supports the idea that α-gal can be secreted in the saliva of *Am. americanum*. Together with basophil activation data, these results suggest the potential role of *Am. americanum* saliva antigens as the primary cause of the delayed-type hypersensitivity reaction, although this requires further investigation.

N-glycan profiling of unfed and partially-fed salivary glands and saliva extracted from *Am. americanum* and *Ix. scapularis* revealed the presence of N-linked glycans with terminal α-gal caps; however, they were absent in *Am. maculatum*. The fact that α-gal is present in the saliva, even in trace amounts, supports the idea that ticks play a role in the induction of a hypersensitivity reaction in humans. Humans without α-gal hypersensitivity are known to have as much as 1% of their circulating IgG antibodies that are specific for anti-gal^36^, which means that the immune system can already recognize this carbohydrate antigen. Because ticks have the ability to remain attached to their host for a prolonged period of time, it is conceivable that small amounts of α-gal and other antigenic molecules being continually secreted into the host could be recognized and initiate an immune response, which resulted in the development of an IgE response directed against α-gal.

Alpha-gal epitopes are commonly expressed on cells and tissues of non-primate mammals, but xylose is found almost exclusively in plants. While we are unsure as to the source, these core-modified glycoforms were found consistently in multiple tissue types and under various conditions, as well as in multiple species. Xylose and core-3-linked fucose may be the most common carbohydrate epitopes recognized by human IgE antibodies^37^. In the literature, β(1,2)-xylose linked to a core mannose has been described in the N-glycans of major pollen allergens, as well as a major peanut allergen^38^.

Mass spectrometry of gel excisions (from Figure 4A) revealed the presence of many proteins and glycoproteins (Supplementary Data Table S1). The protein database for *Ix. scapularis* is well populated and contains many sequences, but the database for *Am. americanum* is scant. The presence of laminin γ-1 was found in both *Ix. scapularis* and *Am. americanum* salivary glands, which has previously been reported to contain an α- gal moiety on the protein, and is suspected of being a common IgE-reactive protein in beef allergy patients in Japan^12^. The alpha chain of type IV collagen was discovered to contain α-gal moieties^39^, and it was identified in our unfed *Ix. scapularis* salivary glands. However, a feature of these proteins is that they lack a signal peptide usually associated with secretion and are therefore not likely to be the primary instigators of the human host α-gal sIgE response. An alternative method by which these proteins may be secreted into the saliva could be via exosomes. Purification of exosomes from tick saliva for the identification of α-gal-containing antigens should be carried out in future studies.

We discovered numerous proteins from both *Am. americanum* and *Ix. scapularis* that could potentially be involved in the α-gal hypersensitivity conundrum. We present a narrowed down list of protein candidates in Table 1, and among these discovered many proteins and enzymes involved in carbohydrate metabolism. Multiple glycoside hydrolases in the salivary glands, which could aid the tick in cleavage of its own and host carbohydrates, are attractive candidate molecules. Unfed *Ix. scapularis* contained the enzyme α-D-galactosidase (EC 3.2.1.22), a signal peptide-containing enzyme responsible for hydrolyzing terminal α-galactosyl residues from glycoproteins and glycolipids that can potentially cleave galactose molecules resulting in free galactose for use in galactosylation elsewhere. Interestingly, we discovered in the lower molecular weight region of *Ix. scapularis* unfed salivary glands, a galactose-binding lectin (galectin) (Uniprot B7Q1V4), and from the higher molecular weight region, a hemolectin. Previously, a group in Japan reported a galectin that can bind galactose containing moieties in *Ornithodoros moubata* ticks^40^. We found a hemelipoglycoprotein and a heme lipoprotein in *Ix. scapularis*, and heme lipoprotein precursors were also found in *Am. americanum*. A considerable amount of research has also been conducted on hemelipoglycoprotein, which has strong binding specificity towards galactose in the tick *Dermacentor marginatus*^41^. Because lectins are present in the salivary glands, it is possible that they are capable of capturing glycoproteins from the host blood. Conceivably, capture of host glycans by tick lectins and cleavage by glycoside hydrolases in conjunction with tick and bacterial glycosyltransferases could result in the α-gal glycan.

We have yet to identify α-1,3-galactosyltransferase in *Am. americanum* or *Ix. scapularis*, but our combined results provide evidence that terminal α-1,3-galactose residues exist in the saliva and salivary glands after the initiation of feeding, both from human and animal hosts, which leads us to believe that there are three possible scenarios that could lead to synthesis:

1. The glycans are captured by lectins and modified with glycoside hydrolases and glycosyltransferases,
2. galactosyltransferase enzymes from bacterial species contained in the normal microbiota or vectored by of *Am. americanum* or *Ix. scapularis* are responsible for the α-gal glycan,
3. some ticks contain an uncharacterized enzyme with a similar or equivalent function to α-1,3-galactosyltransferase, which is yet to be investigated.

Immunoblotting, immunolocalization, and glycan profiling demonstrated that α-gal exists in the salivary proteins of *Am. americanum* and *Ix. scapularis* but not *Am. maculatum.* These data, in tandem with the significant upregulation of CD63+ expression on human basophils indicating activation after inoculation with *Am. americanum* and *Ix. scapularis* salivary antigens, but not *Am. maculatum* (Figure 7), strengthens the idea that bites from α-gal-containing/producing ticks could be involved with the onset of AGS. This represents a significant step forward in our understanding of the sensitization of humans to carbohydrates by ticks, and the clinical implications of tick bites in the United States and worldwide.

The results described in this study provide new insight into tick physiology and support the possibility of hypersensitivity reactions instigated after parasitism by ticks. This research helps to further our understanding of the process in which *Am. americanum* and *Ix. scapularis* obtain and transmit pathogenic α-gal to the host. Our hope is that this mechanism can be used in the future to treat or protect humans from a plethora of medical conditions. This study also highlights the need for allergists and clinicians to consider *Ix. scapularis* and *Am. americanum* bites when diagnosing red meat allergy cases.

## Authors’ Contributions

Conceived and designed the experiments: SK, SPC, GD.

Performed the experiments: GC, SH, SK.

Analyzed the data: GC SPC, GD, PA, SK.

Contributed new reagents/materials/analysis tools: SK, SPC.

Wrote the paper: GC, SPC, SK.

All authors read and approved the manuscript.

---

**Supplement 1.**
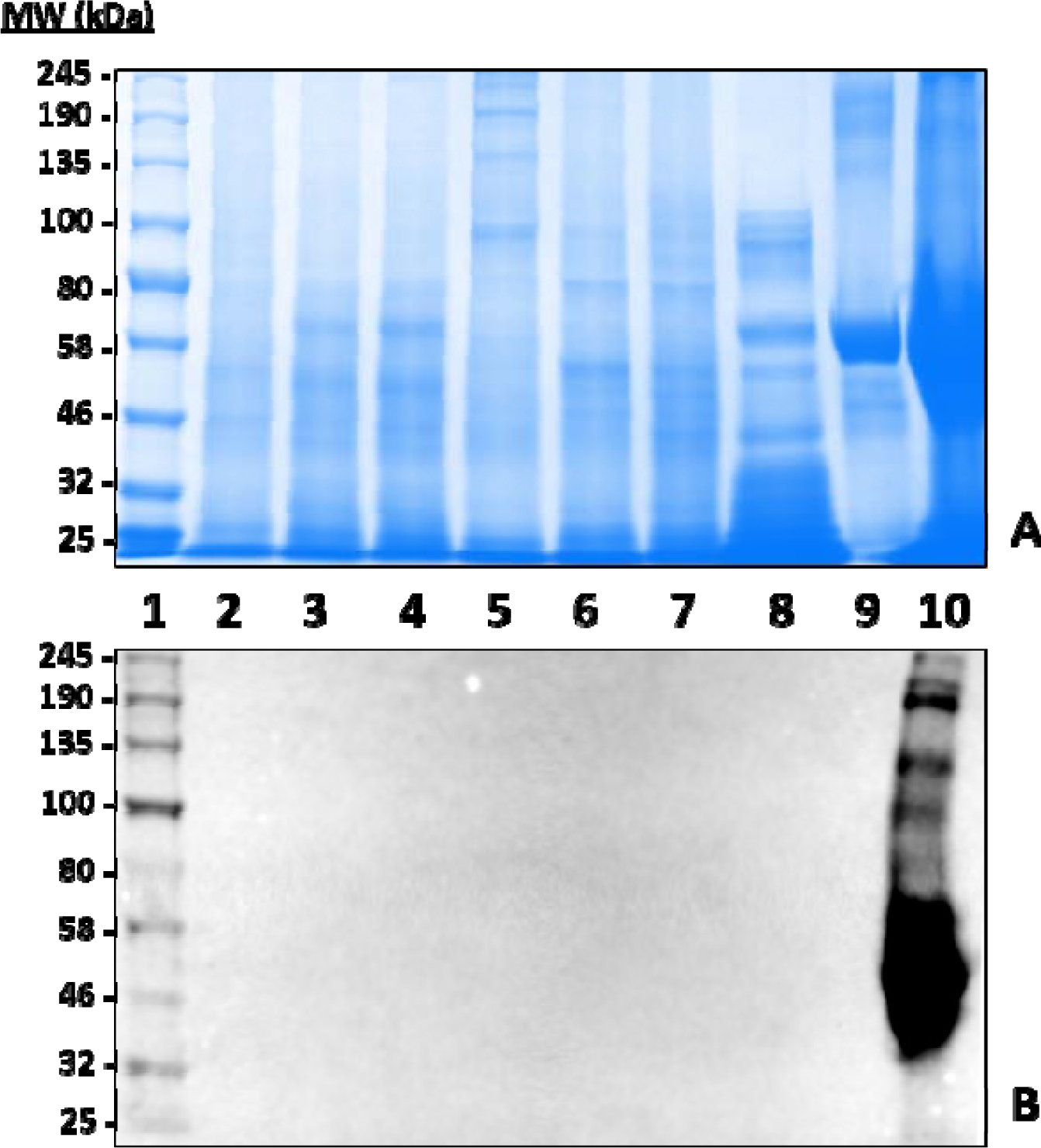
Identification of α-gal-containing antigens in the *Amblyomma maculatum* (*Amac*) midgut (MG) and salivary gland (SG) tissues. A total of 15 µg unfed (UF) and partially-fed (PF) MG and SG tissue homogenates and saliva samples were fractionated on **A)** 12.5% SDS-PAGE, and **B)** western blot using anti-alpha-gal antibody. Lane 1: A broad range (11–245 kDa) pre-stained protein standard, Lane 2: *Amac* UF MG, Lane 3: *Amac* (3 days) PF MG, Lane 4: *Amac* (8 days) PF MG, Lane 5: *Amac* UF SG, Lane 6: *Amac* (3 days) PF SG, Lane 7: *Amac* (8 days) PF SG, Lane 8: *Amac* saliva, Lane 9: Bovine serum albumin and, Lane 10: Diluted whole sheep blood (1:100).

**Supplement 2.**
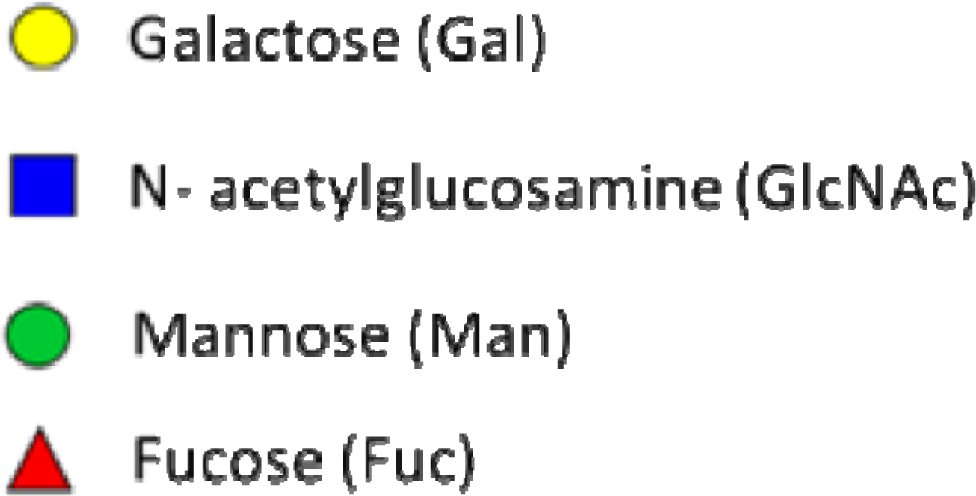
Key of glycan nomenclature used in figures. The above color-filled shapes are a guide to the identification of carbohydrates found in the saliva and salivary glands of *Amblyomma americanum* ticks. Yellow-filled circles are galactose, blue-filled squares are N-acetylglucosamine, green-filled circles are mannose, and red-filled triangles are fucose.

